# Enforced Symmetry: The Necessity of Symmetric Waxing and Waning

**DOI:** 10.1101/328070

**Authors:** Niklas Hohmann

## Abstract

This paper demonstrates that the symmetric waxing and waning patterns that can be observed in ecological measures such as occupancy and geographic range are created by averaging, rescaling, conditioning and combining different sources of information. Therefore symmetric waxing and waning from the origination of a taxon to the extinction of a taxon in any measure should be treated as the null hypothesis and non-informative.

## 1 Introduction

A central question in ecology is whether taxa follow a trajectory from their origination to their extinction, be it in the number of species, the number of individuals or the spatial presence. [1] To derive information about these fundamental processes, measures such as occupancy, which is commonly defined as the proportion of an area in which a taxon can be found in, geographic range, which is often defined as the latitudinal interval in which a species occurs, and species richness are used [2] [3] [4]. Prior research has shown that most taxa display a symmetric pattern in these measures, consisting of an increase in the measure from the origination up to the middle of their life span, followed by a decrease in the measure that is axis symmetric to the increase in the early days of their life span [5]. This pattern is called the symmetric waxing and waning of taxa and can, for example, be observed in occupancy of fossil molluscs [3]. Cases of asymmetric waxing and waning have been observed as well [6], and trajectories of occupancy have been interpreted in different ways. They were, for example, linked to the inability of taxa to occupy niches for longer periods of time [3], the Red Queen Hypothesis [7], or to different types of interaction between taxa that are determined by occupancy [6]. Occupancy and geographic range were further linked to extinction risk [3] [8] [9].

The aim of this paper is to demonstrate that there are mechanisms inherent to the procedure that is used to create the trajectories of taxa that will automatically lead to an increase in symmetry. This is independent of the measure of interest, so any measure that will be analysed with this procedure will display an increased symmetry.

The procedure takes any set of curves describing the measure of interest throughout time. First, each curve is shifted to begin at zero (x-shift), then rescaled to end at one (x-scale). Then the amplitudes of the curves are rescaled (y-scale) and all rescaled curves are averaged [5] [3]. One part of this procedure that will generate symmetry is averaging. This is demonstrated by Donsker’s invariance principle ([10], p. 474), which displays that averaging different random processes can lead to the same, symmetric random process (fig. 1). Another influence is the blurring effect of background noise that is being introduced into the analysis by the x-shift and the x-scale. This combines different sources of noise from different time intervals and of different taxa, which can drown possible signals. The last effect discussed here that increases symmetry is conditioning. This describes the effect that the extinction and the origination of the taxa is already known, which puts an upper and a lower bound on the developments of taxa in their lifespan. This makes strongly asymmetric developments impossible (figs. 2, 3, 4). Because of these three effects, symmetric waxing and waning of any measure of a taxon should not only be treated as the null hypothesis, but as non-informative.

**Figure 1:**
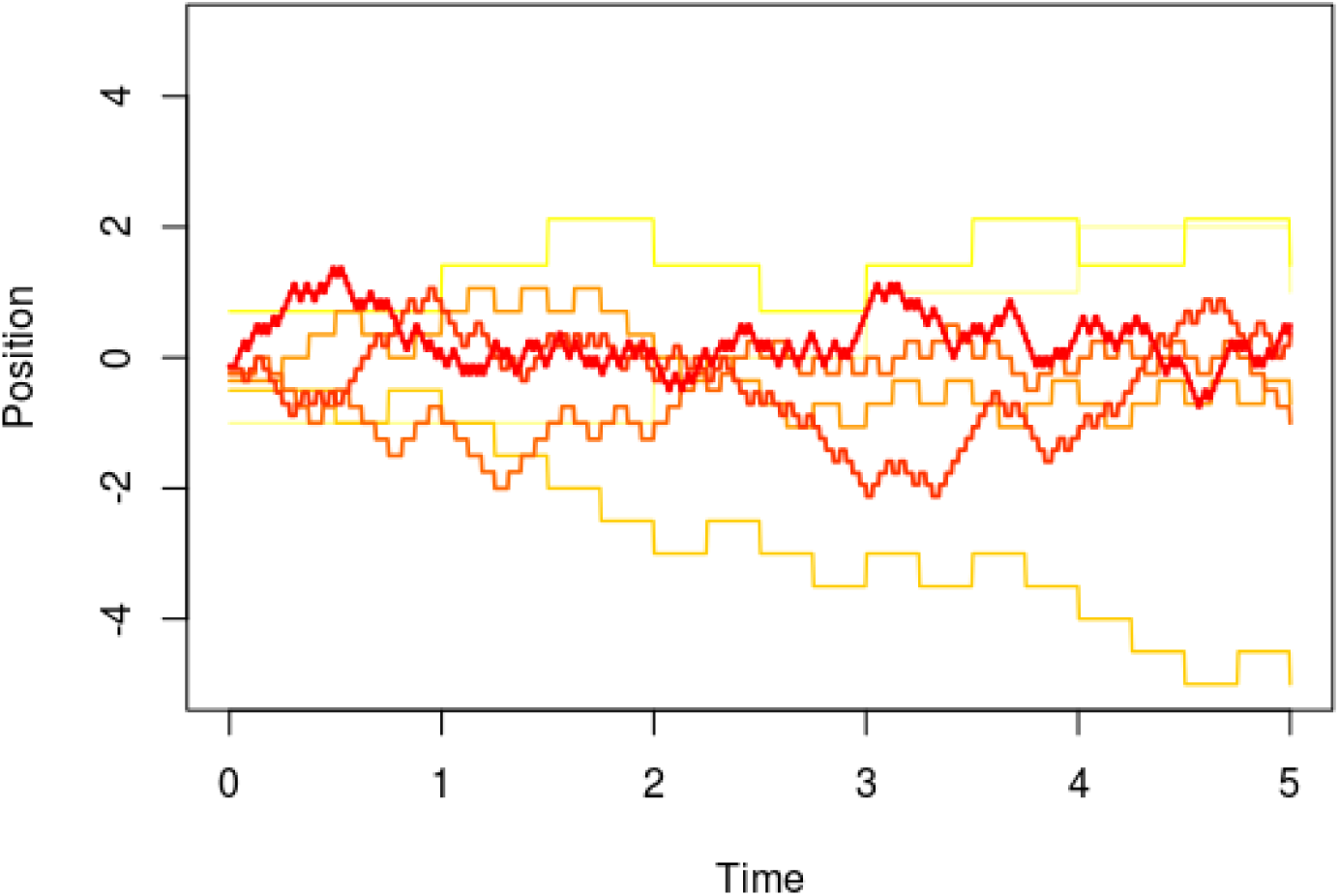
An example for Donsker’s invariance principle as described in [10]. The underlying stochastic process is a symmetric random walk. The more the color turns to red and the thicker the lines get, the better the approximation of the Brownian motion. Approximations are shown for *n* = 1, 2, 4, 8, 16, 32, 64. *n* = 1 corresponds to the thin yellow line, *n* = 64 to the thick red line.

**Figure 2:**
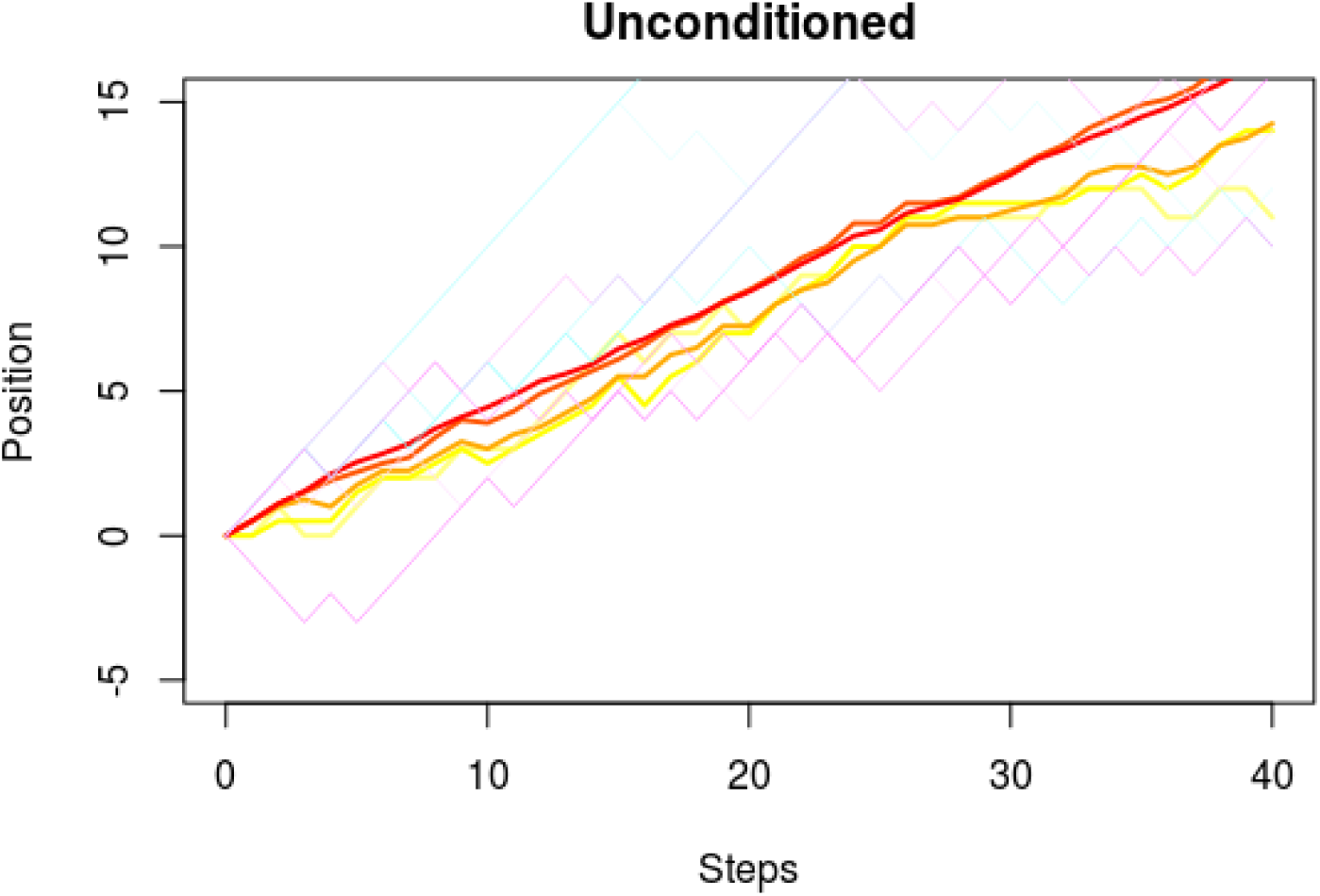
The unconditioned case for an asymmetric walk with transition probability *p* = 0.7. The yellow to red lines are averages of the first *k* paths for *k* = 2, 4, 8, 20, 100. The case *k* = 2 is represented by the light yellow line, the case *k* = 100 by the red line, intermediate values of *k* are orange. Colder colors show eight arbitrary paths of the random walk. The tendency of the random walk to increase is clearly visible.

## 2 Symmetry by Averaging

One way to generate symmetry out of randomness is averaging. This is done in different ways in probability theory. One of the best known ways to achieve symmetry by averaging is the central limit theorem. It roughly states that the averaged sum of random variables converges towards the highly symmetric standard normal distribution ([10], p. 317). This is even the case for the sum of random variables with different distributions ([10], p. 319). Similar results exist for stochastic processes. One such result is Donsker’s invariance principle, which states that averaged sums of stochastic processes will converge to the Brownian motion ([10], p. 474) (fig. 1). Therefore Donsker’s invariance principle can be seen as an infinite dimensional version of the central limit theorem, with the Brownian motion replacing the standard normal distribution. Just like the standard normal distribution, the Brownian motion is highly symmetric:

- It is self-similar in the sense that rescaled versions of a Brownian motion are again a Brownian motion ([10], p. 455)
- It is independent of time reversal in the sense that if (*W*_*t*_)_(*t∈*[0,1])_ is a Brownian motion, then so is (*W*_1_ *− W*_1*−t*_)_(*t∈*[0,1])_.

It is important to note that the invariance under time reversal of the Brownian motion is a reformulation of the axis symmetry as it is for example seen in [5], fig 2. This is since invariance under time reversal means that it is indistinguishable whether the plots where created from origination to extinction or vice versa. But this is equivalent to axis symmetry along an axis that is parallel to the y-axis and shifted by 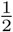 to the right. Similar formulations to this are also valid for real-valued functions. Here the axis symmetry of a function *f* with respect to an axis that parallel to the y-axis but shifted by *a/*2 is equivalent to *f* (*a − x*) = *f* (*x*), which is again invariance under time reversal.

So if the trajectories of any of the measures for which symmetric waxing and waning has been observed are some type of stochastic process then these effects of symmetrization by averaging will appear. It is important to note that this symmetrization will not only occur under the much used null hypothesis of a symmetric random walk1, but also under the much broader class of stochastic processes with expectation value zero and finite variance (see [10], 474 for technical details). Therefore a process that generates symmetry is not necessarily a symmetric random walk.

## 3 Symmetry by Noise

Another effect of rescaling and averaging is that noise from different sources is combined, which can make it impossible to detect signals.

If, for example, occupancy trajectories of different taxa are given, then each trajectory at a given time contains information about different effects on said taxon at this time. These effects can, among others, include changes in environmental conditions or interactions with other taxa. By rescaling these trajectories to start in zero and to end in one, signals that were originally in different time intervals are then present in the same interval between zero and one. Averaging these rescaled trajectories combines all these different signals from different sources and different time intervals on different taxa. This leads to a strong background noise that can drown a potential signal.

To make these considerations more precise, a random walk model is used. For an ordinary asymmetric and homogeneous2 random walk, first a series of i.i.d. (independent identically distributed) random variables (*X*_*i*_)_*i∈*ℕ_ satisfying

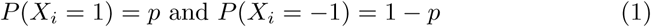

is defined. Here *p* is a value between zero and one. Then the asymmetric homogeneous random walk (*Y*_*n*_)_*n∈*ℕ_ is defined as

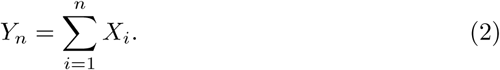

This is the classical symmetric random walk if 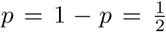. The value of *p* is called the transition probabilitiy, since it determines the probability of the random walk to transition to a higher (and a lower) position.

Distinguishing two potential transition probabilities is important for the question of waxing and waning, since they contain the signal about the tendency of the random walk to increase or decrease. To measure the distinguishability of the transition probabilities of two random walks, the relative entropy is used. This is supported by Stein’s theorem ([11], p. 452), which states that the speed of convergence of type 2 error is determined by the relative entropy. For a random walk with transition probability 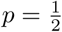 and a random walk with transition probability 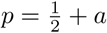, the relative entropy is given by

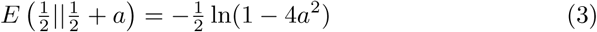

Here, *a* formalizes a signal, given as the deviation from the null hypothesis of the transition probability 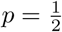.

To evaluate the effects of noise on distinguishability, the noise is modeled by making the transition probabilities random. So for a noise influenced random walk, the transition probabilities are given by

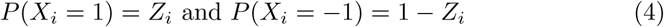

with *Z*_*i*_ being a random variable taking a.s. (almost surely) values in (0, 1). In this case the relative entropy is given by

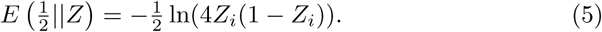

Here, the relative entropy is random just like the transition probabilities. To exactely evaluate the effect of noise, a distribution of the random variable *Z*_*i*_ has to be fixed. If for example the signal is *a* = −0.05 and the noise is a uniform distribution from 0.45 to 0.55, then the relative entropies of the deterministic transition probabilities and the random transition probabilities differ by a factor of six. So in this case it is six times harder to distinguish the noise-influenced signal from the symmetric transition probability than it is to distinguish the signal without the noise from the symmetric transition probability.

Although the extend of this effect is dependent on the type of noise present, this example demonstrates that noise can make the recognition of deviations from a null hypothesis much harder.

## 4 Symmetry by Conditioning

Another effect that increases symmetry is conditioning, which means that there is additional information available that has to be taken into account. In the case of waxing and waning, this additional knowledge is that the origination and extinction are at fixed points. Because of this additional knowledge, some trajectories of taxa are not feasible. Taxa cannot decline too much in the first half of their life span, since otherwise they would go extinct. Neither can they increase too much in the second half of their life span, since they have to go extinct at a fixed point. This leads to a necessary increase in the first half and a neccesary decrease in the second half of a taxons life, which is based on conditioning.

This can be illustrated by the first 2*n* time steps of a random walk which is starting at zero and has a transition probability *p*. Every path of this random walk up to time 2*n* consists of increasing steps and decreasing steps, which are coded by +1 and –1. Overall there are 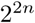 possible paths. If 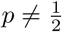, then some paths will be more common than others and some paths are less common. For 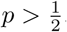, the paths with many increasing steps will be more common, whereas the paths with many decreasing steps will be more rare. In general, paths with a high or low ratio of increasing steps to decreasing steps will be rare or common, whereas paths with a moderate ration of increasing steps to decreasing steps will be moderately frequent. So the asymmetry in the transition probabilities is most clearly expressed by the very rare and the very common paths (see fig. 2 for an example with *p* = 0.7).

For simplicity, only the case of conditioning to a fixed end point is discussed here (fig. 3). The case of only positive paths follows mutatis mutandis (fig. 4). Denote the set of all paths that end at zero by *Z*_*p*_. Then by the law of the elementary conditional probability, the probability of a path conditioned on the fact that it ends at zero is given by

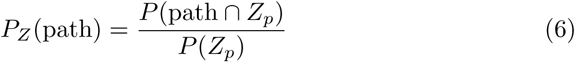

**Figure 3:**
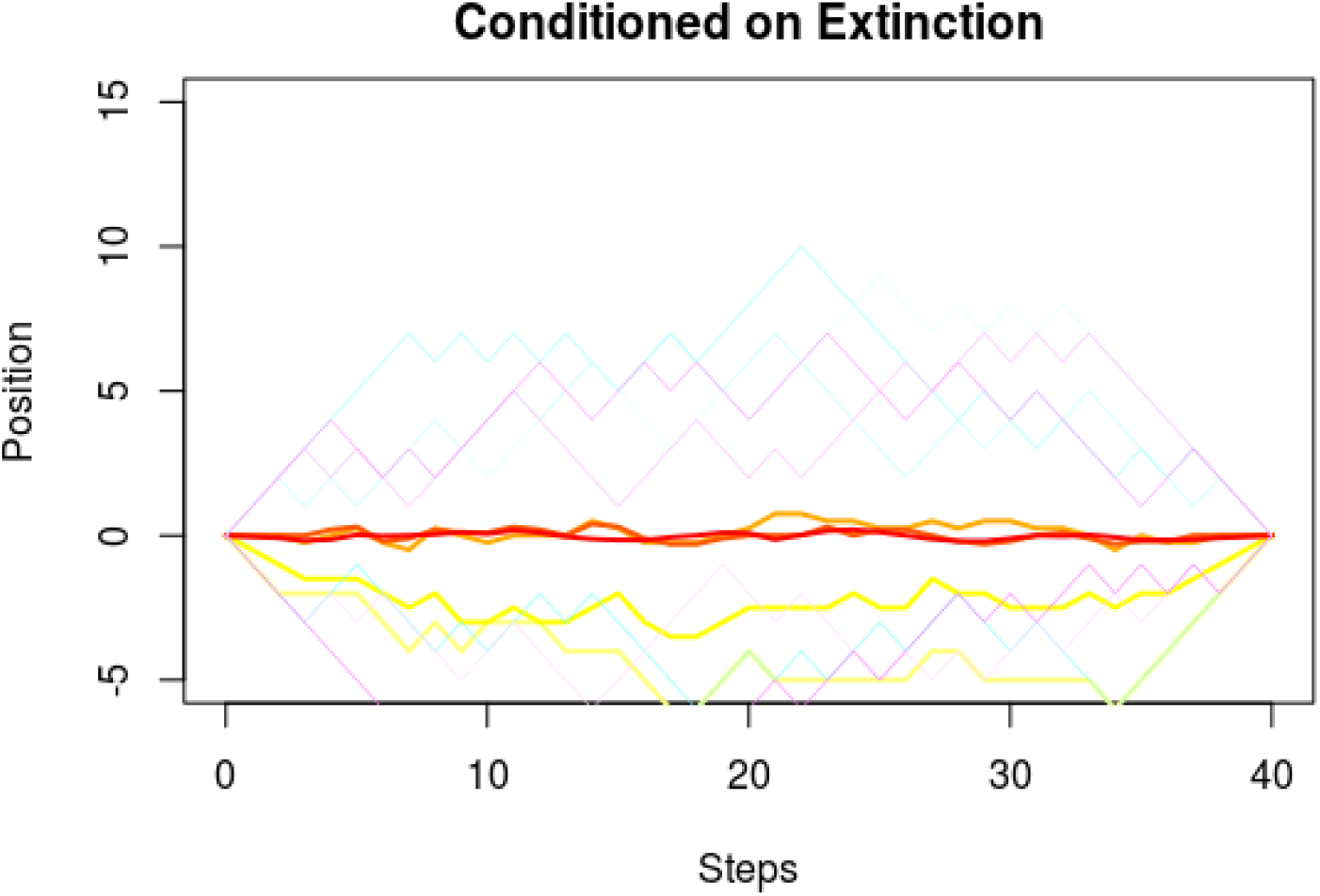
The asymmetric random walk as displayed in fig. 2, but conditioned on hitting zero after 40 steps. Color codes are as in fig. 2. The averaged paths approximate the x-axis, and non-averaged paths display a high degree of axis symmetry.

**Figure 4:**
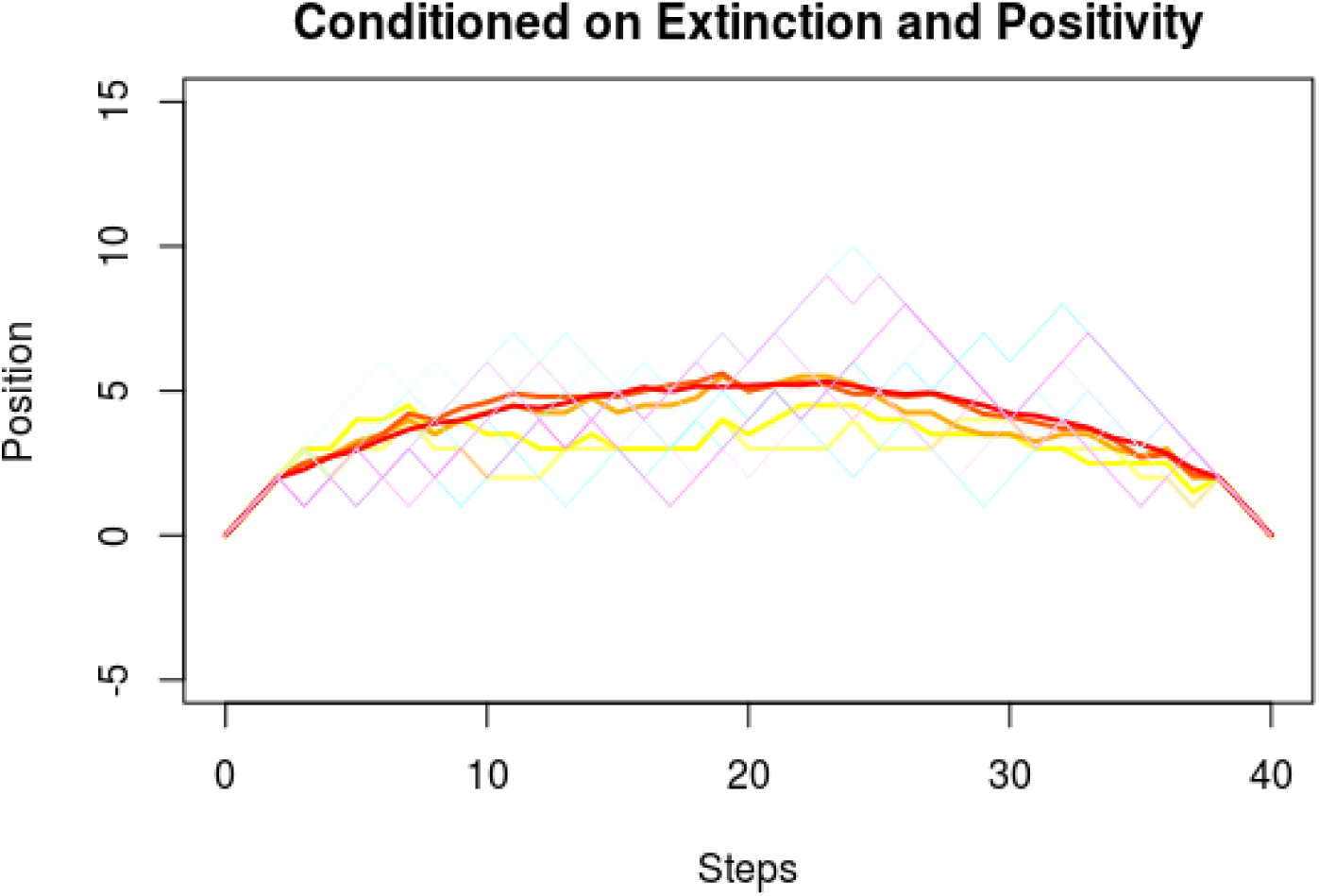
The asymmetric random walk as displayed in fig. 2, but conditioned to be strictly positive and to hit zero after 40 steps. Color codes are as in fig.2. The averaged paths converge to an axis-symmetric curve, the asymmetry of the underlying random walk is not visible any more.

Here *P*_*Z*_ is the probability after conditioning on the set *Z*_*p*_. So every path that does not end at zero is assigned the probability zero and all the probabilities of all paths that end at zero are multiplied with the constant 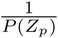.

But as discussed above the paths that will generate most of the asymmetry are the paths with a high or a low ratio of increasing steps to decreasing steps. These paths are all assigned the probability zero, since only paths with a ratio of increasing steps to decreasing steps of one end at zero. So conditioning leads to a focus on paths that have a balanced proportion of increasing steps to decreasing steps, which leaves out all paths that display a lot of asymmetry.

This can be made more formal by embedding the set of all paths {1, −1}^2*n*^ into ℝ^2^*n*, where the paths form the vertices of a hypercube. Then random variables in ℝ^*2n*^ with distributions *P* and *P*_*Z*_ can be used to examine the long term behaviour of averages of the conditioned and unconditioned paths. One approach is the multidimensional central limit theorem ([10], p. 326), which emphasises the importance of the expectation value vector. It corresponds to the center of mass of the hypercube if the vertices are weighted by their probability. By conditioning, vertices with high or low weights are deleted, which leads to a shift of the center of mass towards paths with a moderate ratio of increasing steps to decreasing steps. This corresponds to an increase in the symmetry of the averaged path.

## 5 Discussion

The three lines of argument above show that

1. A large class of random processes will generate symmetric trajectories when they are averaged (section 2)
2. Fixing trajectories to predetermined times of origination and extinction increases symmetry (section 4)
3. Any signal that deviates from symmetry is hard to recognize in the presence of noise (section 3)

There are certainly more effects relevant to the generation of symmetric waxing and waning than the three effects explained above. But the lines of argument discussed here show that there is a tendency inherent to the procedure used to create these trajectories that will always increase symmetry.

### 5.1 Possible Solutions

This raises the question whether there is a way to derive information about the trajectories without using the procedure described in the introduction. The hardest question here is whether there is a way to distinguish between a symmetric waxing and waning that is a statistical artefact and a driven symmetric waxing and waning that is biologically meaningful in the sense that it is generated by an underlying process that is connected to the stage of life of taxa. One approach for random walks could be to examine the lengths of consecutive increasing steps or decreasing steps. Here a random walk that is driven by symmetric waxing and waning (in the sense that it has a high transition probability in the first half of its life span and a low transition probability in the second half of its life span) displays a different pattern that a symmetric random walk (fig. 5), although both have symmetric averaged paths (fig. 6). The driven symmetric waxing and waning has fewer short runs than the waxing and waning generated by the symmetric random walk, but up to 40 times more long runs. This might be a possible approach to derive information about trajectories without averaging.

**Figure 5:**
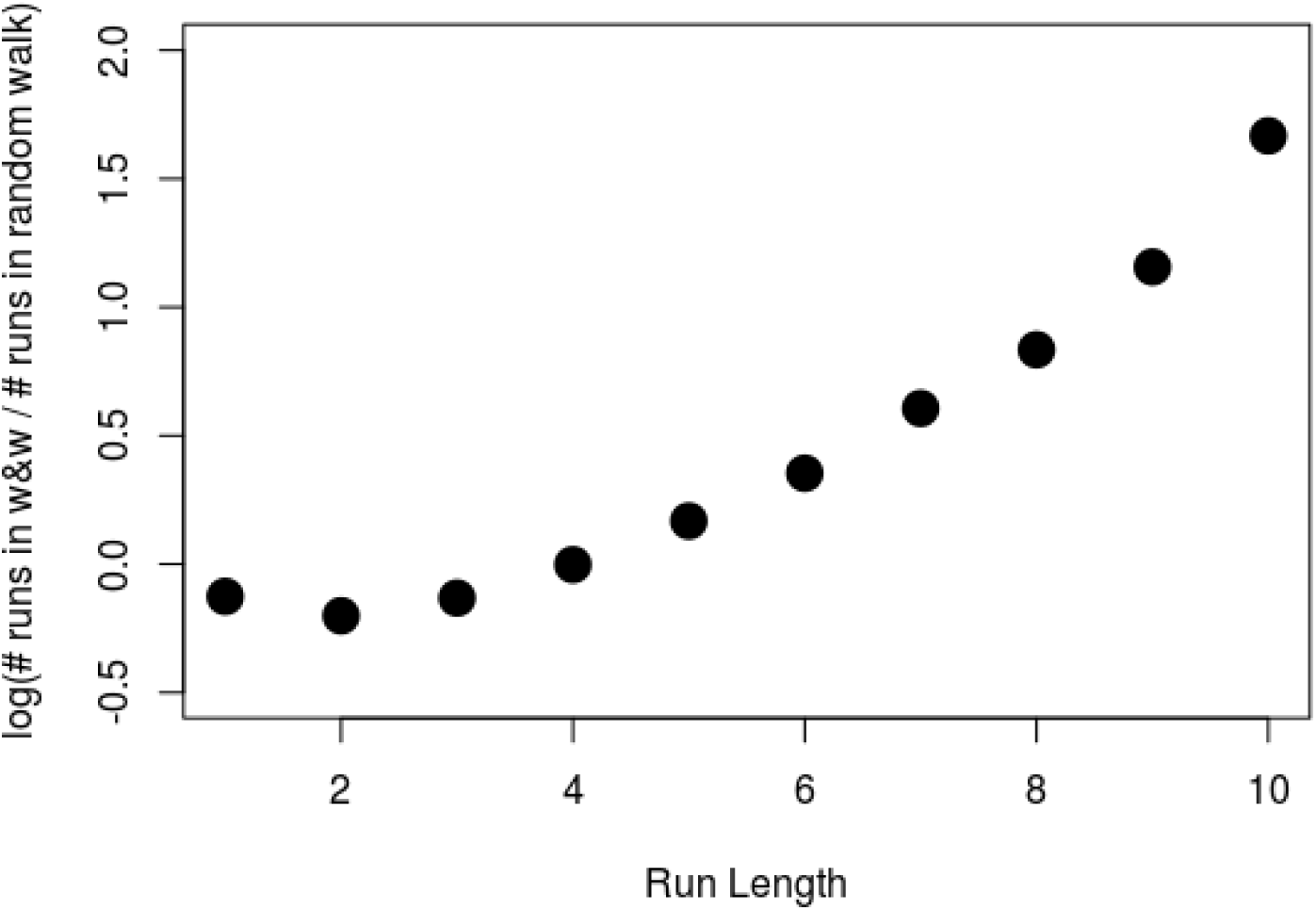
The ratio of the number of runs with a given length from the driven waxing and waning and the symmetric random walk. Both processes are conditioned to be positive and hit zero after 20 steps for the first time. The symmetric random walk has a constant transition probability of *p* = 0.5, the driven symmetric waxing and waning is defined to have a transition probability of *p* = 0.7 for the first ten steps and a transition propability of *p* = 0.3 for the last ten steps. A run is a series of consecutive increases or decreases in the path. Note that the y-axis is logarithmic with base 10. This shows that the symmetric random walk will have more short runs than the driven waxing and waning, whereas the driven waxing and waning has a lot more long runs. This might be an approach to distinguish between an artefactual symmetric waxing and waning, which is here represented by the symmetric random walk, and a driven symmetric waxing and waning.

**Figure 6:**
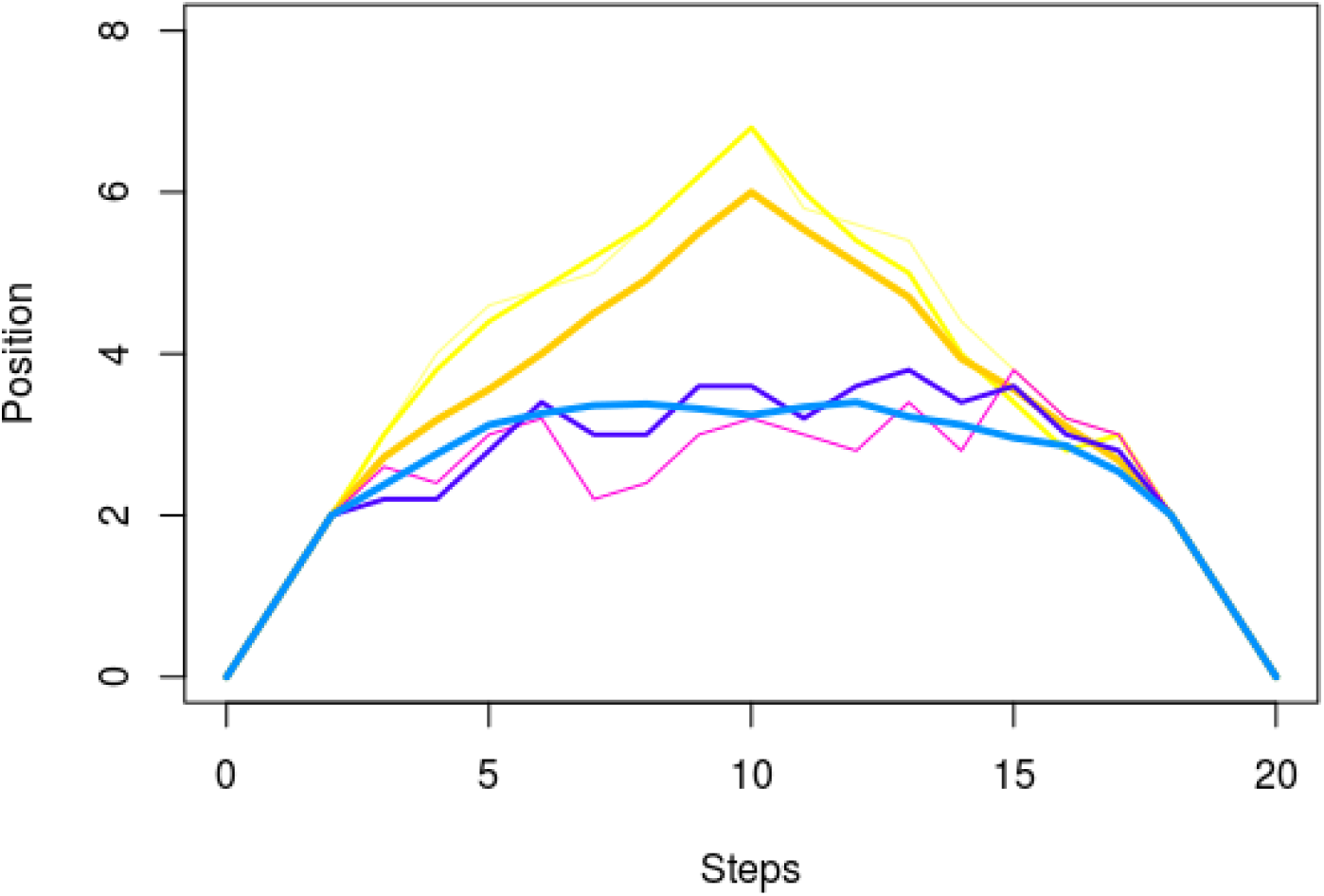
Averaged paths of a symmetric random walk and a driven symmetric waxing and waning as explained in fig. 5. Orange and yellow are averaged paths of the driven symmetric waxing and waning, blue are averaged paths of the symmetric random walk. Thin lines are averages over five paths, medium lines are averages over 10 paths and thick lines are averages over 100 paths.

It is important to remember that most paleoecological data is binned, for example in time intervals or geographic grid cells. This can lead to situations where a symmetric function generates asymmetric bins (fig. 7) or a asymmetric function generates symmetric bins (fig. 8). This effect is weaker for functions spanning multiple bins, therefore taxa found only within few bins should be excluded from the analysis.

**Figure 7:**
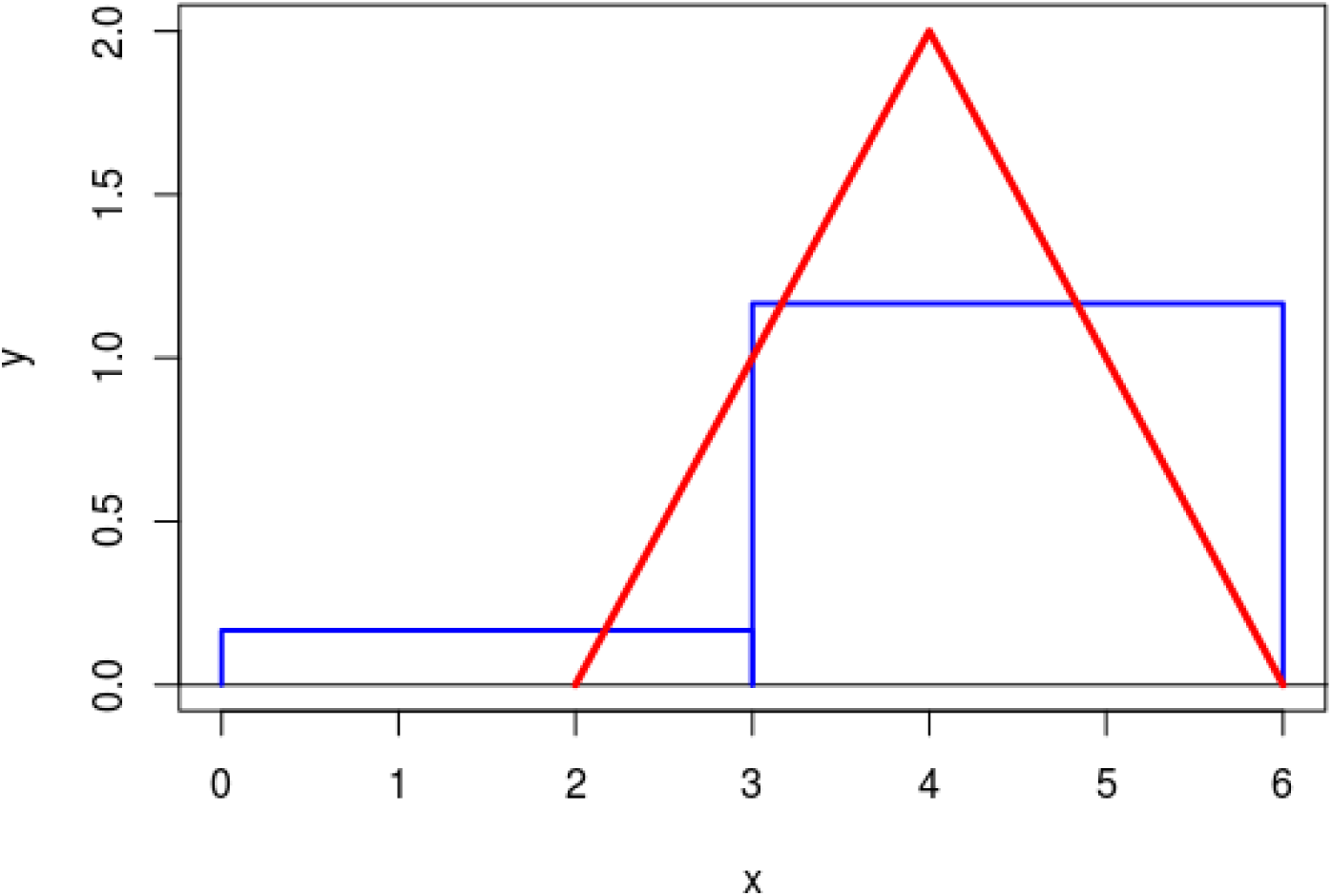
A symmetric function (red), generating asymmetric bins (blue). The area of a bin corresponds to the area under the function in the bin.

**Figure 8:**
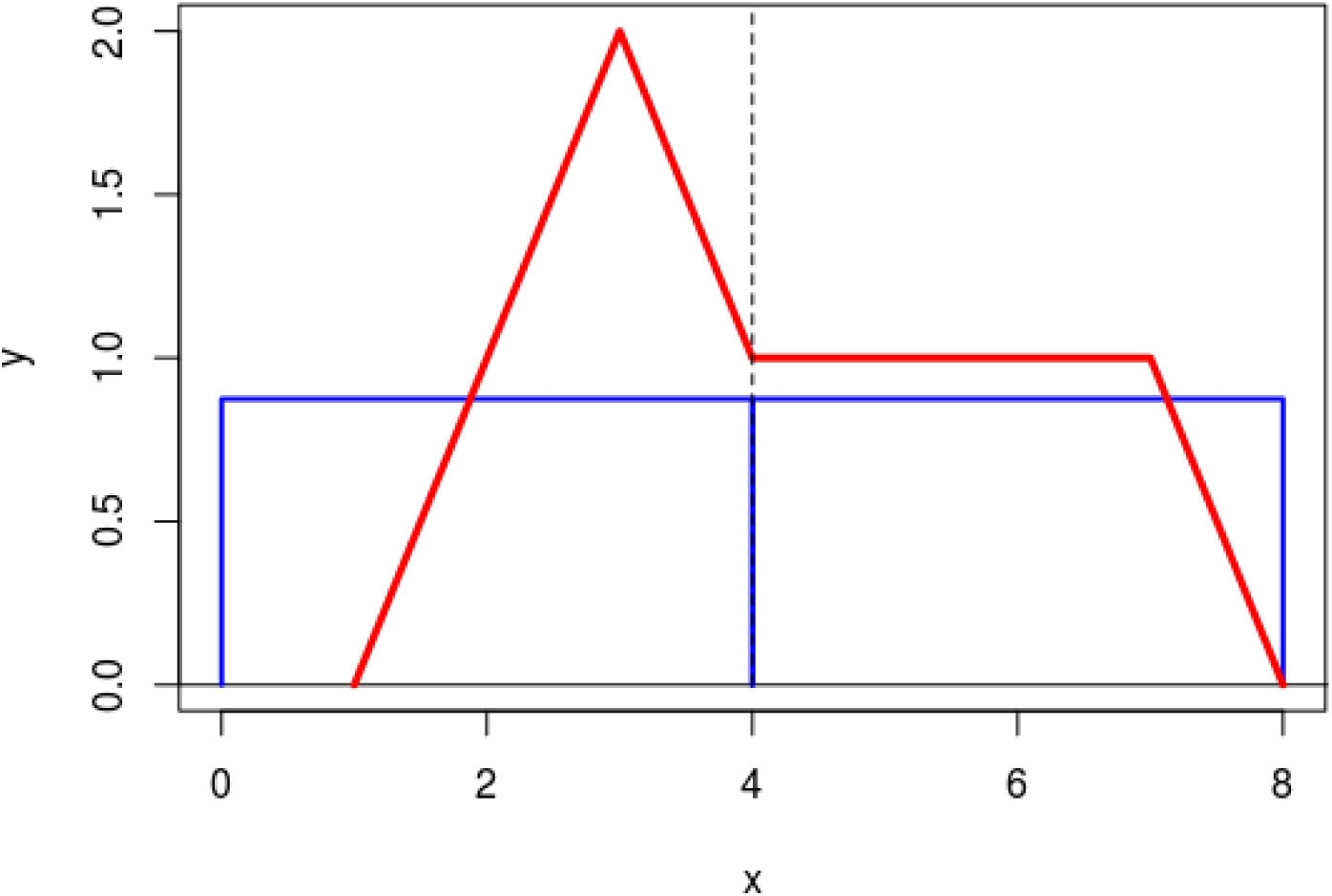
An asymmetric function (red), generating symmetric bins (blue). The area of a bin corresponds to the area under the function in the bin.

### 5.2 Asymmetric waxing and waning

Cases with asymmetric waxing and waning as described in [6] can contain important information. But asymmetry can also be generated by a lack of available data. So in cases where asymmetric waxing and waning appears, it is recommendable to first check whether the data is sufficient to derive any conclusions and then look for possible sources of the asymmetry. Potential sources of asymmetry can then be examined and isolated in the raw data with the help of more specific statistical approaches.

## 6 Summary

- If trajectories are averaged, prone to noise, or conditioned on their origination and extinction, then some type of symmetric waxing and waning will always appear. Therefore symmetric waxing and waning should be treated as the null hypothesis and as non-informative
- For trajectories that display asymmetric waxing and waning, it is important to observe possible sources that can generate asymmetry and examine them separately
- To derive information about trajectories of taxa, methods that do not rely on averaging and conditioning should be used. Looking at run lengths or behaviour of paths under permutations might be possible approaches to distinguish hypotheses about trajectories of taxa

## 7 Acknowledgements

I would like to thank Wolfgang Kiessling for pointing me towards the problem and discussing the topic as well as giving important input. Further thanks go to my supervisor Emilia Jarochowska for her incredible patience as well as constant feedback and encouragement. Thanks also go to the YouTube channel Numberphile for a continuous supply with amazing videos about math, most notably the espisode about the hot hand featuring Lisa Goldberg. It served as inspiration for the proposed solution described above.

## Footnotes

[1 For a strict definition of symmetric random walk see below equation (2)

[2 In the sense of having constant transition probabilities over time

